# Chromosome-length haplotigs for yak and cattle from trio binning assembly of an F1 hybrid

**DOI:** 10.1101/737171

**Authors:** Edward S. Rice, Sergey Koren, Arang Rhie, Michael P. Heaton, Theodore S. Kalbfleisch, Timothy Hardy, Peter H. Hackett, Derek M. Bickhart, Benjamin D. Rosen, Brian Vander Ley, Nicholas W. Maurer, Richard E. Green, Adam M. Phillippy, Jessica L. Petersen, Timothy P. L. Smith

## Abstract

**Background:** Assemblies of diploid genomes are generally unphased, pseudo-haploid representations that do not correctly reconstruct the two parental haplotypes present in the individual sequenced. Instead, the assembly alternates between parental haplotypes and may contain duplications in regions where the parental haplotypes are sufficiently different. Trio binning is an approach to genome assembly that uses short reads from both parents to classify long reads from the offspring according to maternal or paternal haplotype origin, and is thus helped rather than impeded by heterozygosity. Using this approach, it is possible to derive two assemblies from an individual, accurately representing both parental contributions in their entirety with higher continuity and accuracy than is possible with other methods.

**Results:** We used trio binning to assemble reference genomes for two species from a single individual using an interspecies cross of yak (*Bos grunniens*) and cattle (*Bos taurus*). The high heterozygosity inherent to interspecies hybrids allowed us to confidently assign >99% of long reads from the F1 offspring to parental bins using unique k-mers from parental short reads. Both the maternal (yak) and paternal (cattle) assemblies contain over one third of the acrocentric chromosomes, including the two largest chromosomes, in single haplotigs.

**Conclusions:** These haplotigs are the first vertebrate chromosome arms to be assembled gap-free and fully phased, and the first time assemblies for two species have been created from a single individual. Both assemblies are the most continuous currently available for non-model vertebrates.

## Background

New technologies and algorithms for chromosome-scale genome assembly have improved the contiguity of reference genomes in the past several years [1]. These new methods are more efficient than previous methods, allowing high-quality assemblies of the genomes of a wider variety of organisms, rather than for model organisms only. In addition to increasing assembly efficiency, these technologies have focused on addressing two of the foremost challenges of genome assembly: long repetitive regions and heterozygosity of diploid genomes. Repetitive regions are difficult to assemble due to their low sequence complexity, resulting in gaps in reference genomes [2,3]. Mitigating this issue, advances in long-read sequencing technologies [4,5] have facilitated the generation of reads longer than many of these repetitive regions, spanning what otherwise would be assembly gaps [6,7].

Advances in sequencing technology have thus far not been as successful at resolving heterozygous regions of diploid genomes as they have been at resolving repetitive regions. Heterozygous loci, especially those containing complex structural differences between the haplotypes, add intractable complexity to the assembly graphs used to assemble genomes. Most current long-read genome assemblers, such as canu [8], flye [9], and miniasm [10], choose a random haplotype in each heterozygous region and save the unused haplotype as an alternate, resulting in a single pseudo-haploid assembly containing sequence from both parental haplotypes. Another long-read assembler, FALCON-unzip, uses long reads spanning multiple heterozygous regions to phase the assembly graph as much as possible, but the assemblies it generates still contain numerous haplotype switch errors [11]. The long-range information present in proximity ligation and linked read libraries has also been used to phase diploid assembly graphs with mixed results [12,13].

Trio binning is a new assembly technique that avoids the need for such complex strategies by deconvoluting the problem of diploid genome assembly into a pair of simpler haploid assemblies [14]. Trio binning uses variation present in short reads from two parents to sort long reads from their offspring into bins representing either maternal or paternal haplotypes. The long reads in these bins are then assembled independently of one another, resulting in two haploid assemblies of higher quality and contiguity than would be possible with a diploid assembly. This method’s ability to correctly infer haplotype of origin for long reads from the offspring is dependent on how divergent the two parental genomes are, as greater divergence results in more places in their offspring’s genome where the two haplotypes are differentiable. Thus, trio binning produced better results for assembly of an intraspecies hybrid of two breeds of cattle (heterozygosity ~0.9%) than for a human trio (heterozygosity ~0.1%) [14].

Here, we apply trio binning to an interspecies F1 hybrid of yak (*Bos grunniens*) and cattle (*Bos taurus*), two species that diverged ~4.9Mya [15] but are capable of producing fertile offspring [16]. The interspecies application of trio binning maximizes the use of heterozygosity to make it easier to bin reads resulting in high-quality reference genomes for both parental species. The resulting fully phased haploid assemblies of both the cattle and yak genomes contain chromosome-arm length haplotigs, representing the most contiguous assemblies to date of large diploid genomes.

## Results

We applied trio binning to a trio consisting of a yak cow (*Bos grunniens*) Molly, a Highland bull (*Bos taurus*) Duke, and their F1 hybrid offspring Esperanza **(Figure 1)**. After verifying Esperanza’s parentage **(Supplementary Table S1)**, we sequenced both parents with Illumina short reads and their offspring with PacBio long reads. We estimated Esperanza’s heterozygosity to be ~1.2%, compared to ~0.9% for the cross-breed cattle hybrid assembled by Koren et al. [14], which is consistent with the longer divergence time between yaks and cattle than between indicine and taurine cattle **(Supplementary Figure S1)**.

**Figure 1.**
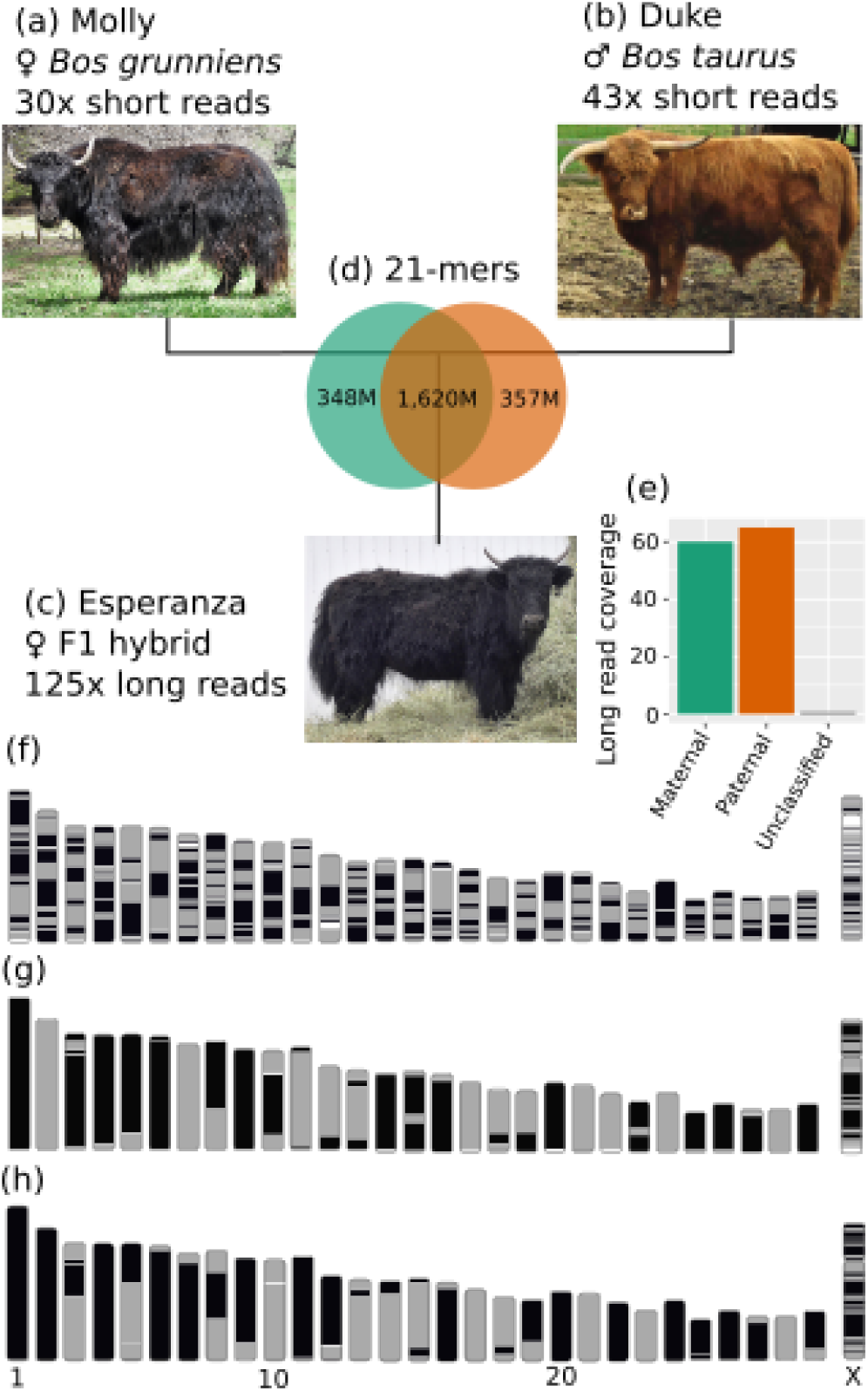
Trio binning of a yak/cattle hybrid. (a-c) We collected short reads from a female yak and a male cattle, and long reads from their F1 hybrid offspring. (d) Counts of 21-mers shared by Molly and Duke and those unique to a single parent. (e) Long-read coverage of the maternal and paternal haplotypes after binning reads from Esperanza using 21-mers from (d). (f-g) Ideograms of contigs on chromosomes for (f) ARS_UCD1.2, (g) Esperanza’s maternal (yak) haplotype assembly, and (h) Esperanza’s paternal (cattle) haplotype assembly, with contigs represented as solid blocks of a single color.

Using the short reads from the two parents, we found ~350 million 21-mers unique to each parental line. More than 99% of the total length of the long reads from Esperanza contained one or more 21-mers unique to one of the parental genomes, allowing them to be sorted into maternal or paternal bins **(Figure 1d,e)**, each of which were then independently assembled.

The initial contig assemblies of these two haplotypes are ultra-continuous **(Figure 1f-h)**, with contig N50s of 70.9 Mb for the yak haplotype and 71.7 Mb for the cattle haplotype. In addition, over one third of the autosomal chromosomes in both assemblies are comprised of a single contig: 15 in maternal and 12 in paternal out of 29. BUSCO [17] analyses of both genomes show most single-copy orthologs present in the initial contig assemblies. 97.1% of single-copy orthologs are present in the maternal assembly, 95.5% of which are both complete and single-copy. 96.8% of single-copy orthologs are present in the paternal assembly, 95.6% of which are both complete and single-copy.

Trio binning assembly is advantageous not only because removing heterozygous diploidy as a complicating factor leads to more contiguous assemblies, but because it results in two fully phased assemblies. To confirm that the maternal (yak) assembly and the paternal (cattle) assembly were correctly phased, with no switch errors, we again took advantage of the large divergence between the two haplotypes resulting from the interspecies cross by testing the similarity of both assemblies to several cattle and yak individuals **(Figure 2 and Supplementary Figures S2 & S3)**.

**Figure 2.**
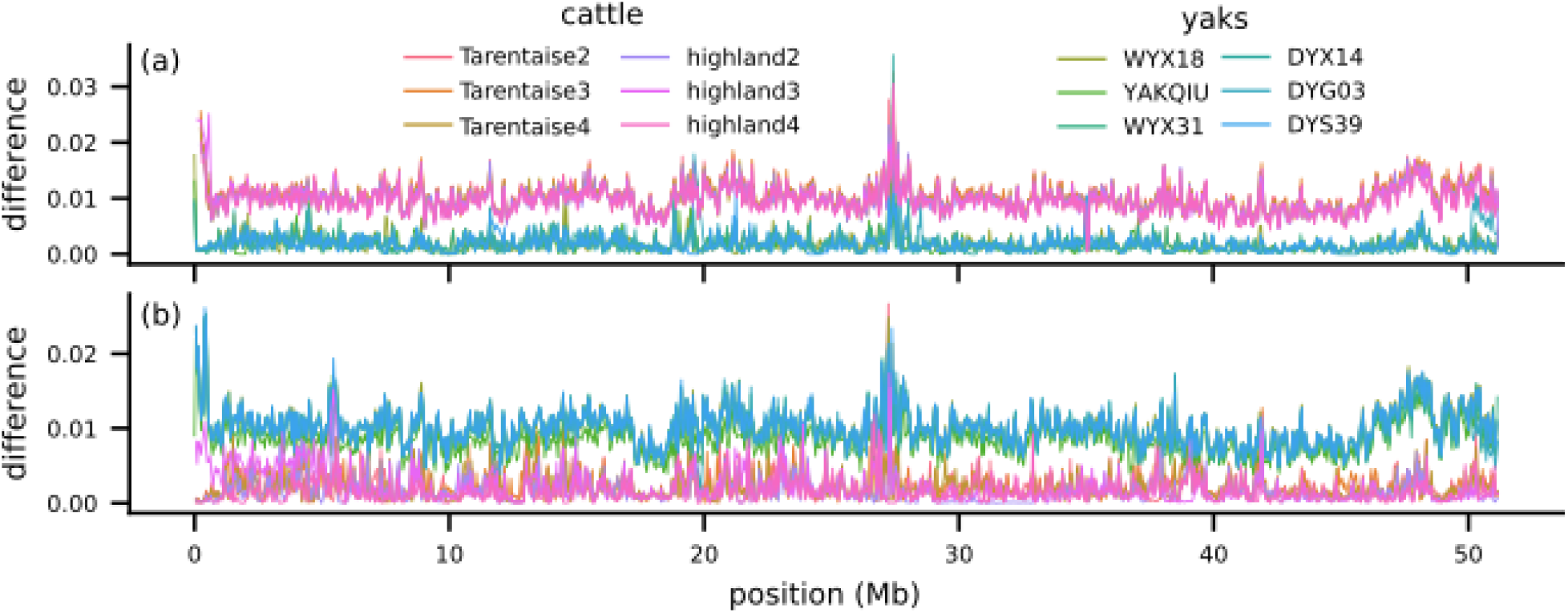
Alignment of six cattle and six yaks to chr29 of our (a) maternal and (b) paternal assemblies shows that the maternal haplotype assembly is more similar to yak genomes than cattle and the paternal haplotype assembly is more similar to cattle genomes, demonstrating that they are phased correctly.

We aligned short reads from three Highland cattle, three Tarentaise cattle, two wild Asian yaks, and four domestic Asian yaks to the paternal and maternal assemblies, and calculated the number of SNPs for each individual compared to both references in 50kb windows across the genome. In a vast majority of windows, the mean SNP rate of the six cattle is higher than that of the six yaks when compared to the maternal yak reference (98.4%), and the mean SNP rate of the six yaks is higher than that of the six cattle when compared to the paternal cattle reference (99.7%). Notable exceptions to this occur in places like the beginning of maternal chr11, where all six yaks have higher SNP rates compared to the maternal reference than all six cattle, indicating that the maternal reference is more cattle-like at these locations. However, the paternal reference is not more similar to the six yaks at the same locations, indicating that these are not likely to be haplotype switch errors. Rather, we hypothesize that these are regions of cattle introgression into the maternal genome, as introgression among various *Bos* species including cattle and yak is known to be pervasive worldwide [18,19].

Some chromosomes in both genomes are comprised of multiple contigs, so scaffolding the assemblies was still necessary. To this end, we sequenced 250 million reads from a Hi-C library created from a tissue sample of Esperanza. The short read length of a Hi-C short read library presents fewer chances in each read for finding kmers unique to one parent, so we instead aligned all read pairs to both the maternal and paternal haplotype assembly and used alignment scores to bin read pairs. We were able to assign 152M Hi-C pairs to one or the other haplotype using this method, and used the remaining 98M pairs to scaffold both assemblies. The resulting scaffolds had an N50 of 86.2Mb for the paternal and 94.7Mb for the maternal assembly.

Both scaffolded assemblies are highly concordant with the current cattle reference genome **(Supplementary Figures S4 & S5)**. Whole genome alignment of the two assemblies to ARS_UCD1.2 revealed a small number of large (>1Mb) structural differences between ARS_UCD1.2 and the yak and cattle haplotypes: four in the yak and five in the cattle haplotype. Further investigation of these discordant segments using a recombination map of cattle [20], an optical map [21], Hi-C heatmaps, the location of telomeric repeats, short read coverage around the breakpoints, and the previous cattle reference UMD3.1 [22], provided sufficient evidence to justify inverting three contigs in the maternal assembly and three contigs in the paternal assembly.

After assigning scaffolds to chromosomes using the recombination map for autosomes and alignment to ARS_UCD1.2 for the X chromosome, we filled gaps created between contigs during scaffolding and chromosome assignment by aligning binned long-reads back to their assemblies using the PBJelly pipeline. This process was able to fill 74 of these gaps in the maternal and 78 in the paternal haplotype assembly, increasing the contig N50s to 79.8Mb for the maternal and 72.8Mb for the paternal assembly. We then finalized both assemblies with a polishing step.

Out of 402 identified gaps on the ARS-UCD1.2 reference assembly, our maternal and paternal assemblies conclusively closed 213 and 219 gaps, respectively (Supplementary Table S3). Gap closure was confirmed by the alignment of 500bp of sequence flanking ARS-UCD1.2 gaps to each assembly and ensuring that the alignments were on the same scaffold, within 100 kb of each other. Gap flanking sequence could not be placed on the same scaffold (trans-scaffold) in 185 and 179 cases for the maternal and paternal assemblies, respectively, suggesting that the cause of ARS-UCD1.2 gaps could be due to scaffolding errors. Of these trans-scaffold closures, 77 and 110 events in the maternal and paternal assemblies were not consistently closed, suggesting structural differences between the assemblies that may indicate true differences between species or individuals.

Intersection of repetitive element annotations with gap flanking sequence revealed that most ARS-UCD1.2 gap regions may have been caused by discrepancies in scaffolding of contigs that were terminated by L1 LINE elements (Supplementary Table S4). These events were followed closely by BovB repetitive elements, which may have also terminated a large proportion of contig ends. While the association of repetitive elements in gap flanking sequence points towards a potential cause for the gap region in ARS-UCD1.2, we cannot rule out the possibility that transposition of L1 LINEs, BovB and other active retroelements may have been spuriously detected in this analysis. Inconsistency of flanking elements around gaps in the sire and dam assemblies (Supplementary Table S5) suggests that several elements may have moved locations after the divergence between *Bos taurus* and *Bos grunniens*.

The final assemblies of both the cattle and yak genome contain the largest contigs and the fewest gaps of any current assembly of a large diploid genome **(Figure 3)**. Both cover the largest two chromosome arms, the q-arms of chr1 (158Mb) and chr2 (136Mb), with a single contig. The maternal yak assembly has 19 gaps on autosomes and 13 gaps on the X chromosome; the paternal highland cattle assembly has 18 gaps on autosomes and 22 gaps on the X chromosome. For comparison, the current cattle reference ARS_UCD1.2 has 260 gaps on autosomes and 55 on the X chromosome; both assemblies reduce this number of gaps by nearly a factor of ten. Furthermore, our trio assemblies of yak and cattle are comparable or superior to other vertebrate reference genomes in terms of contig N50, number of gaps, and size of largest contig compared to size of largest chromosome arm.

**Figure 3.**
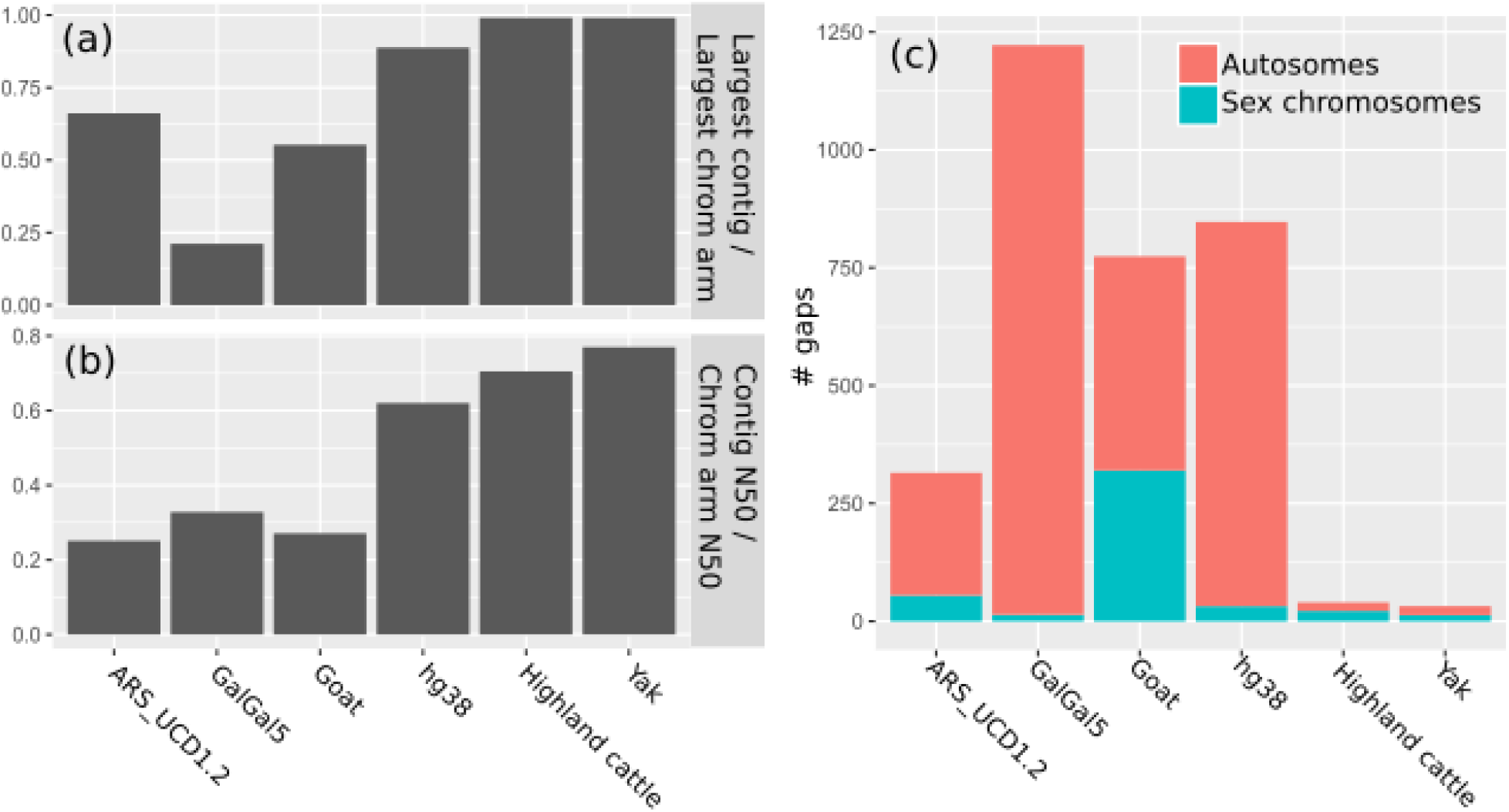
Comparison of trio highland and yak assemblies to current cattle, chicken, goat, and human reference assemblies, based on ratio of largest contig size to largest chromosome arm size (a), ratio of contig N50 to chromosome arm N50 (b), and number of gaps in autosomes and the major sex chromosome, i.e., X in cattle, yak, goat, and human and Z in chicken (c). We note that the number of gaps in hg38 is somewhat inflated due to its gapped assembly of centromeres.

Trio binning also resolves heterogeneous loci into haploid sequences. BOLA, the bovine major histocompatibility complex, is a set of highly diverse loci on chr23 containing variants associated with infectious disease susceptibility [23,24], The trio assemblies of the cattle and yak haplotype both contain all four subclasses of BOLA in a single contig. The coat-color gene KIT is another example of a heterogeneous locus that these assemblies resolve into ungapped true haploid sequences for both haplotypes [25,26].

## Discussion

The application of trio binning to a yak/cattle hybrid trio demonstrates that this method is capable of producing highly accurate reference assemblies more continuous than those currently available for species with large diploid genomes. The initial contig N50s of the maternal yak and paternal cattle assemblies, at 70.9Mb and 71.7 Mb, respectively, are larger than the contig N50s of the current references for yak (20.4kb) [15] and cattle (ARS_UCD1.2, 25.9Mb) [27]. Our assemblies are also more continuous than the previous trio binning assemblies of bovines, at 23.3Mb and 26.6Mb for the maternal and paternal haplotypes of an Angus × Brahman cross [14]. Thus, our initial haplotig assemblies, even before scaffolding and gap-filling, represent large improvements over existing assemblies of the cattle and yak genomes.

These assemblies not only represent large improvements compared to the current cattle reference genome, but are more contiguous by some measures than even the highest-quality reference genomes of organisms such as human (hg38), chicken (galgal5), and goat [28]. For example, the largest contig in hg38 is a 132Mb contig containing most of the 140Mb q-arm of chr4, whereas over one third of the q-arms of the all-acrocentric autosomes in our assemblies are comprised of a single contig.

Moreover, these assemblies used as input only long reads from a single individual and short reads from its parents, including the mitochondrial genomes, which were assembled from parental short reads. We also used a Hi-C library to scaffold the assemblies and various orthogonal data types to correct errors in the scaffolding and assign scaffolds to chromosomes, but many chromosomes in both haplotypes were assembled into single contigs in the initial long read assembly and thus did not require these additional data types. By comparison, recent chromosome-scale assemblies of other non-model mammals such as horse [29] and goat [28] required many additional data types, such as Sanger sequence, BAC clones, Chicago libraries, optical maps, and linked reads, to achieve their levels of contiguity and composition. We used a pre-existing genetic map for validation of our assemblies, but the high contiguity and accuracy of our scaffolded but otherwise unedited contig assemblies demonstrates that long reads plus a Hi-C library are sufficient for producing high-quality assemblies using trio binning.

Furthermore, our results demonstrate that it is now technologically feasible to assemble full chromosome arms gap-free with only long reads. The remaining gaps in our assembly are likely the result of repetitive regions such as rDNA, centromeres, and large segmental duplications too large to be spanned by the long reads we have, but the ever-increasing maximum read lengths achievable with SMRT [4] and nanopore [5] sequencing continue to surpass the sizes of new repetitive regions. We predict that these improvements to existing technologies, along with algorithmic advances such as those that enabled assembly of the human Y centromere [30], will therefore make gap-free assemblies of vertebrate genomes possible in the near future.

These assemblies are not only highly contiguous, but have the additional advantage of being fully haploid rather than pseudo-haploid as in most current reference assemblies of large diploid genomes. This is especially valuable in highly heterogeneous regions of the genome where the two haplotypes in an individual are most likely to be divergent. We show that the haploid assemblies produced by trio binning can fully resolve difficult to assemble heterogeneous loci such as MHC without the need for additional phasing data. This technique is likely to represent a large benefit in the assembly of out-bred or wild vertebrate species that are known to produce viable hybrids.

Trio binning using a cross-species hybrid, in addition to allowing for easier binning of long reads through increased heterozygosity, also has the advantage of producing reference genomes for two species with long reads from only a single individual. Thus, this approach will be especially useful for comparative genomics studies in which contiguous haploid reference genomes for two related species can be used to identify evolutionary breakpoints with high accuracy.

## Conclusions

Our assembly of chromosome-length haplotigs for the yak and cattle genomes using trio binning suggests that trio binning is the best approach currently available for assembling the genomes of diploid organisms that either can be cross-bred with a closely related species or at least have enough population structure within the species to allow breeding two unrelated parents with divergent genomes. While many organisms of biological interest are polyploid or unable to be bred in a controlled setting, many model organisms and other highly studied species would be good candidates for trio binning. We expect that this method will therefore soon be used to assemble new reference genomes for a variety of species.

## Methods

### Sample collection and preparation

All animal protocols were approved by the Institutional Animal Care and Use Committee of the University of Nebraska-Lincoln, an AAALAC International Accredited institution (IACUC Project ID 1648). Whole blood (EDTA) was collected via jugular venipuncture from the Highland bull and yak cow. Tissue sampling of the yaklander heifer was conducted after euthanization using pentobarbital administered intravenously (1ml/101b). Lung tissue was flash frozen and stored at −80°C until DNA isolation and sequencing.

### Long-read library preparation and sequencing

Genomic DNA was extracted from Esperanza lung tissue using high salt extraction method as described by [28]. The DNA was converted into sequencing libraries using the SMRTbell Express Template Prep Kit (Pacific Biosciences, Menlo Park CA) as directed, except without any shearing step. Three libraries were prepared, one with a 25 kb cutoff setting on the BluePippin instrument (Sage Science, Beverly MA) and two with a 30 kb cutoff setting. The libraries were sequenced with 44 cells on a Sequel instrument using Sequel Sequencing kit v2.1 chemistry (Pacific Biosciences, Menlo Park CA).

### Short-read library preparation and sequencing

Genomic DNA from Esperanza lung (used also for long read sequencing, above) was converted into sequencing libraries using the TruSeq DNA PCR-Free LT Library Preparation Kit (Illumina Inc., San Diego CA) as directed. The shearing was conducted on a Covaris S220 instrument (Covaris Inc., Woburn MA) with setting to 350 base pair fragment size. The same procedure was used to create libraries for parental and unrelated yak samples, except the DNA was prepared from blood using a standard phenol:chloroform extraction as described [31]. Sequencing was performed by 2×150 base paired end sequencing on a NextSeq500 instrument (Illumina Inc., San Diego CA) using High Output Kit v2 (300 cycles) kits.

### Hi-C library preparation and sequencing

33.36 mg frozen lung from Esperanza was removed from cold storage and homogenized by chopping with a sterile scalpel. The resulting lung paste was transferred to a microcentrifuge tube along with one milliliter PBS. Paraformaldehyde (EMS Cat. No. 15714) was added to a final concentration of three percent and the sample was vortexed briefly before rotation for twenty minutes at room temperature. Collagenase from a Dovetail Hi-C Library Preparation Kit (Catalog No. 21004) was added to the crosslinked tissue and incubated for 1 hour at 37°C in an agitating thermal mixer. The liquid phase was taken from this reaction and brought to a final concentration of one percent SDS.

Crosslinked chromatin was bound to SPRI beads and washed thoroughly before digesting with DpnII (20 U, NEB Catalog No. R0543S) for 1 hour at 37°C in an agitating thermal mixer. Biotin-11-dCTP (ChemCyte Catalog No. CC-6002-1) was incorporated by DNA Polymerase I, Klenow Fragment (10 U, NEB Catalog No. M0210L) for thirty minutes at 25°C. Following another wash, intra-aggregate ligation with T4 DNA Ligase (4000 U, NEB Catalog No. 0202T) was carried out overnight at 16°C. Crosslinks were reversed in an eight percent SDS solution with Proteinase K (30 g, Qiagen Catalog No. 19133) for fifteen minutes at 55°C followed by forty-five minutes at 68°C. After SPRI bead purification, DNA was split into two replicates and sonicated to an average length of 350 bp using a Diagenode Bioruptor NGS platform.

Sheared DNA samples were run through the NEBNext Ultra II DNA Library Prep Kit for Illumina (Catalog No. E7645S) End Preparation, Adaptor Ligation with custom Y-adaptors, and SPRI bead purification steps before Biotin enrichment via Dynabeads MyOne Streptavidin C1 beads (ThermoFisher Catalog No. 65002). Indexing PCR was performed on streptavidin beads using KAPA HiFi HotStart ReadyMix (Catalog No. KK2602) and subsequently size selected with SPRI beads.

We sequenced 250M reads of this library on a 2×151bp run of an Illumina NextSeq500 using High Output Kit v2 (300 cycle) kits (Illumina Inc., San Diego CA).

### RNA sequencing

Total RNA was extracted from tissues using Trizol reagent (Thermofisher Scientific) as directed. The RNA-seq method for quantitative transcript abundance estimation was performed by preparation of KAPA RNA HyperPrep Kit stranded RNA libraries as recommended by the manufacturer (KAPA Biosystems Inc., Woburn MA). Each library was sequenced to a minimum of 30 million clusters (paired reads) on a NextSeq500 instrument using 2×150 paired end sequencing as described above. The libraries averaged approximately 300 base inserts and were sequenced with longer-than-standard read lengths, to improve the ability to assign transcripts to the parental genome of origin by detection of parent-specific kmers in the reads.

### Heterozygosity estimation

Heterozygosity of Esperanza was estimated using Genomescope [32].

### Parentage confirmation

Using the Illumina whole genome shotgun sequence data generated for this project as well as other data already published to the public domain, we calculated the number of sites relative to the bovine reference genome that did not follow the expected pattern of inheritance. For example if the sire was homozygous for allele A, the dam homozygous for allele B, and the progeny homozygous for B, in the absence of a genotyping error, this pattern suggests that the reported sire is not in fact the sire. We expect some genotyping errors [33,34], but whatever exclusions are identified when analyzing the verifiable trio should be dwarfed in number when one of the actual parents is swapped in the analysis with an unrelated animal. For this comparison we did trio analysis of the yak × cattle offspring versus the reported Highland sire and yak dam as well as the reported dam versus four unrelated Highland bulls, and the reported sire versus an unrelated yak dam.

The UnifiedGenotyper [35] was used in gt_mode=DISCOVERY to analyze the mapped datasets (bam files) for cattle x yak progeny in turn vs. a prospective sire/dam pair to identify sites polymorphic in the trio, then genotype those positions producing a vcf file. A custom java program was written to search the dataset for exclusions. Given the nature of this cross it was expected that the majority of the sites identified would be those specific to the interspecies mating. Specifically, we ignored all polymorphic sites with the interspecies cross signature of the bovine sire homozygous for an allele A (consistent with the bovine reference allele), the yak dam homozygous for allele B (likely consistent with the allele fixed in yak) and the progeny heterozygous A/B. Since this pattern would be common to any cattle × yak mating, it would not be suitably specific for a parentage test. The sequence data from the animals not part of the trio were generated for another study with a much lower fold coverage requirement. The coverage for these other animals is on average ~14×. It has been demonstrated previously (cattle and sheep genotyping papers) that a genotyping accuracy of ~98% can be attained at this level of coverage. The ~2% error rate in that work was attributable to an undersampling of the second allele for heterozygous genotypes or allele dropout in the assay based genotyping platform. This will have the effect of increasing the rate of exclusions in those animals not reported to be the parents, but it should be at a rate of approximately 1% of the total genotypes analyzed. This error amounts to a small contribution to the observed exclusion count for the negative controls. The results are shown in **Supplementary Table S1**. The reported yak dam produced ~12 fold fewer exclusions than the negative control dam (4.99% vs 0.42%) and the reported Highland sire produced ~32 fold fewer exclusions when compared with the negative control sires (6.24%,5.70%,5.76%, and 7.06%) vs 0.19%). These results indicate a correct parental assignment.

### Contig assembly

The trioBinning scripts from (https://github.com/skoren/triobinningScripts) were used to classify the reads. Briefly, meryl from canu 1.7.1 was used to count all parental k-mers. The k-mers specific to both the maternal and paternal haplotype were identified via the meryl difference command. Finally, any paternal k-mer occurring at least 6 times and any maternal k-mer occurring at least 4 times were retained for classification:

meryl -B -C -m 31 -s maternalIlluma.fa -o mom -threads 28 -memory 60000
meryl -B -C -m 31 -s patemallIlluma.fa -o dad -threads 28 -memory 60000
meryl -M difference -s dad -s mom -o dad.only
meryl -M difference -s mom -s dad -o mom.only
meryl -Dt -n 6 -s dad.only |awk ‘{if (match($1,”>“)) { COUNT=substr($1, 2, length($1)); } else {print $1” “COUNT}}’ |awk ‘{if ($NF < 100) print $0}’ > dad.counts
meryl -Dt -n 4 -s mom.only |awk ‘{if (match($1,”>“)) { COUNT=substr($1, 2, length($1)); } else {print $1” “COUNT}}’ |awk ‘{if ($NF < 100) print $0}’ > mom.counts

Reads with no parental marker were not used in downstream analysis. Classified reads were assembled with Canu 1.7.1 with the patch for truncated consensus (git commit e42d54d4f1b1133b8e944b09733806bfe63bc600) command ‘genomesize=2.8g’ ‘correctedErrorRate=0.105’ ‘cnsErrorRate=0.15’ ‘corMhapSensitivity=normal’ ‘ovlMerThreshold=500’.

### Phasing confirmation

We confirmed that our assemblies are phased correctly by comparing both references to several yak and cattle genomes. We downloaded short reads from SRA for three Highland cattle, three Tarentaise cattle, four domestic yaks, and two wild yaks. Supplementary Table S2 lists the IDs and SRA accessions of these individuals. We aligned short reads to both maternal and paternal haplotype assemblies using bwa mem v0.7 [36] with default parameters and sorted alignments and removed PCR duplicates using samtools sort and rmdup [37] with default parameters. Finally, we called SNPs and calculated window SNP rates, which we define as (# homozygous SNPs + 0.5 * # heterozygous SNPs) / (# bases genotyped in window), using samtools mpileup output piped to a custom script available at https://github.com/esrice/misc-tools/blob/master/pileup2windows.py. We used the mpileup parameters “-Q 20 -q 20” to exclude low-quality base calls or alignments from the pileup, and we did not call SNPs for positions where the sequencing depth was below the 2.5th percentile or above the 97.5th percentile position-depth for that sample.

### Scaffolding

We preprocessed the Hi-C reads by trimming to the DpnII junction sequence GATCGATC. To separate the junction-split Hi-C read pairs into maternal and paternal bins, we aligned all reads to both maternal and paternal contig assemblies using bwa mem v0.7 [36] with default parameters. We then ran the classify_by_alignment program (https://github.com/esrice/trio_binning v0.2.1) to determine based on the ‘AS’ tag of the resulting bam files whether each read pair aligned better to the maternal contigs, the paternal contigs, or both equally. If the read pair aligned better to one haplotype than the other, we used it to scaffold only this haplotype, but if it aligned equally well to both, we used it to scaffold both haplotypes. We then ran SALSA2 v2.2 [38] to scaffold both assemblies using the parameters ‘-e GATC -m yes’.

### Quality control

To find possible mis-assemblies in the scaffolds, we aligned them to ARS_UCD1.2 [27] using mashmap v2.0 [39] with parameter ‘--perc_identity 95’. We also aligned probes from a recombination map of cattle [20] to the scaffolds using bwa mem v0.7 [36]. We examined resulting alignments for each chromosome for evidence of disagreements between our assembly and ARS_UCD1.2 or the recombination map. Where such disagreements existed, we used the combination of evidence from Hi-C heatmaps, ARS_UCD1.2, the recombination map [20], an optical map [21], telomeric repeat location, the previous cattle reference UMD3.1 [22], and short read coverage around the breakpoint to determine whether there was sufficient evidence to edit our assembly to better match the reference. In total, we inverted the orientation of three haplotigs in the paternal assembly and three haplotigs in the maternal assembly.

### Chromosome assignment

We used the alignments of recombination map probe sequences as described above to order and orient scaffolds onto chromosomes. As the recombination map does not include the X chromosome, we used the mashmap alignments between our assemblies and ARS_UCD1.2 to order and orient scaffolds onto the X chromosome.

### Gap filling

We filled remaining gaps in each assembly using the PBJelly pipeline [40], which we modified for compatibility with current versions of the software upon which it depends: blasr [41] v5.3.2 and networkx [42] v2.2. This modified pipeline is available at https://github.com/esrice/PBJelly.

### Gap analysis

Gap flanking sequence consisting of 500 bp of sequence from the 5’ and 3’ ends of each gap region was extracted from the ARS-UCD1.2 reference genome. These flanking sequences were aligned to the sire and dam haplotig assemblies using bwa mem v0.7 [36] and checked for consistency. If both gap flanking sequences were on the same scaffold, were within 100 kb [43] distance of each other, and had no intersecting gaps from the same assembly, the gap was considered closed. Repetitive elements were identified using RepeatMasker (http://repeatmasker.org), with the settings “-q”, “-species cow” and “-no_is.” Repeat annotations were converted to bed coordinates and were intersected with gap flanking regions using Bedtools [43].

### Polishing

Arrow from SMRTanalysis v5.1.0.26412 (pbcommand v0.6.7, arrow 2.2.2 ConsensusCore v1.0.2, ConsensusCore2 v3.0.0, pbalign version: 0.3.1) was used via the ArrowGrid pipeline (https://github.com/skoren/ArrowGrid). Only classified reads were used to polish each haplotype. Initial contigs were polished with two rounds of Arrow. Final gap-filled assemblies were again polished with two rounds of Arrow, using SMRTanalysis v6.0.0.47841 (pbcommand v1.1.1, arrow 2.2.2, ConsensusCore v1.0.7, ConsensusCore2 v3.0.0, pbalign version: 0.3.1).

## List of abbreviations

Mb: megabases
kb: kilobases
MYA: millions of years ago
MHC: major histocompatibility complex
SMRT: single molecule real time

## Declarations

### Ethics approval and consent to participate

All animal procedures were approved by the Institutional Animal Care and Use Committee of the University of Nebraska - Lincoln (UNL, Protocol 1648). UNL is an AAALAC International accredited institution.

### Consent for publication

Not applicable.

### Availability of data and material

The datasets generated and/or analysed during the current study are available in the NCBI BioProject repository under accessions PRJNA551500 and PRJNA552915.

### Competing interests

The authors declare that they have no competing interests.

### Funding

This work was supported by ARS Project Number 3040-32000-034-00D. ESR and sample collection costs were supported by funding from an Enhanced Research Collaboration grant from the University of Nebraska-Lincoln, Institute of Agriculture and Natural Resources, Agricultural Research Division and the USDA Meat Animal Research Center. DMB was supported by USDA CRIS project number: 5090-31000-026-00-D. SK, AR, and AMP were supported by the Intramural Research Program of the National Human Genome Research Institute, National Institutes of Health.

### Authors’ contributions

This study was conceived and designed by TPLS with input from JLP and AMP. TH and PH identified the animals and collected ante-mortem samples; BVL, MPH, and JLP were responsible for handling the euthanasia and tissue collection postmortem. TPLS was responsible for long and short read genomic and RNA sequencing, except NWM and ESR prepared Hi-C sequencing libraries. ESR, AR and SK generated the assemblies. ESR, SK, DMB, and BDR conducted quality control and assessment of the assemblies. TSK performed additional computational analyses. MPH, BVL, and REG provided additional input. ESR, SK, AMP, JLP, and TPLS wrote the manuscript. All authors read and approved the final manuscript.

## Acknowledgements

We thank Beth Nelson and Nedda F. Saremi (UC Santa Cruz), Kristen Kuhn, Kelsey McClure, Jacky Carnahan, and Bob Lee (USMARC) for technical assistance, and Sarah Rada-Scott for providing animals, tissue samples, and photography. Tissue sampling was conducted with help from the following: David Steffen, John Kammermann, Shauna Tietze, Kristin Beede, Erin Duffy, Rachel Burrack, Brent Johnson, Matthew Quinn, Shawna Clement, Lianna Walker, Kylee Sutton, Hiruni Wijesena, Lizzy Dorwart, Katie Howser, Robert Posont, Kerri Bochantin, Taylor Barnes, Lauren Kett, Rebecca Swanson, Joslyn Beard, Aaron Cowell, Emma Winters, Devon Lockman, Jaden Carlson, Madeline Pelster, and Adam Bassett (University of Nebraska-Lincoln); and (Nebraska Wesleyan University) Leah Treffer. This work used the computational resources of the NIH HPC Biowulf cluster (https://hpc.nih.gov), the USDA CERES cluster (a part of the SCINet initiative: https://www.ars.usda.gov/scinet/), and the University of Nebraska - Lincoln Holland Computing Center (https://hcc.unl.edu/). Mention of trade names or commercial products is solely for the purpose of providing specific information and does not imply recommendation or endorsement by the U.S. Department of Agriculture. USDA is an equal opportunity provider and employer.

## Supplementary Figures

**Supplementary Figure S1.**
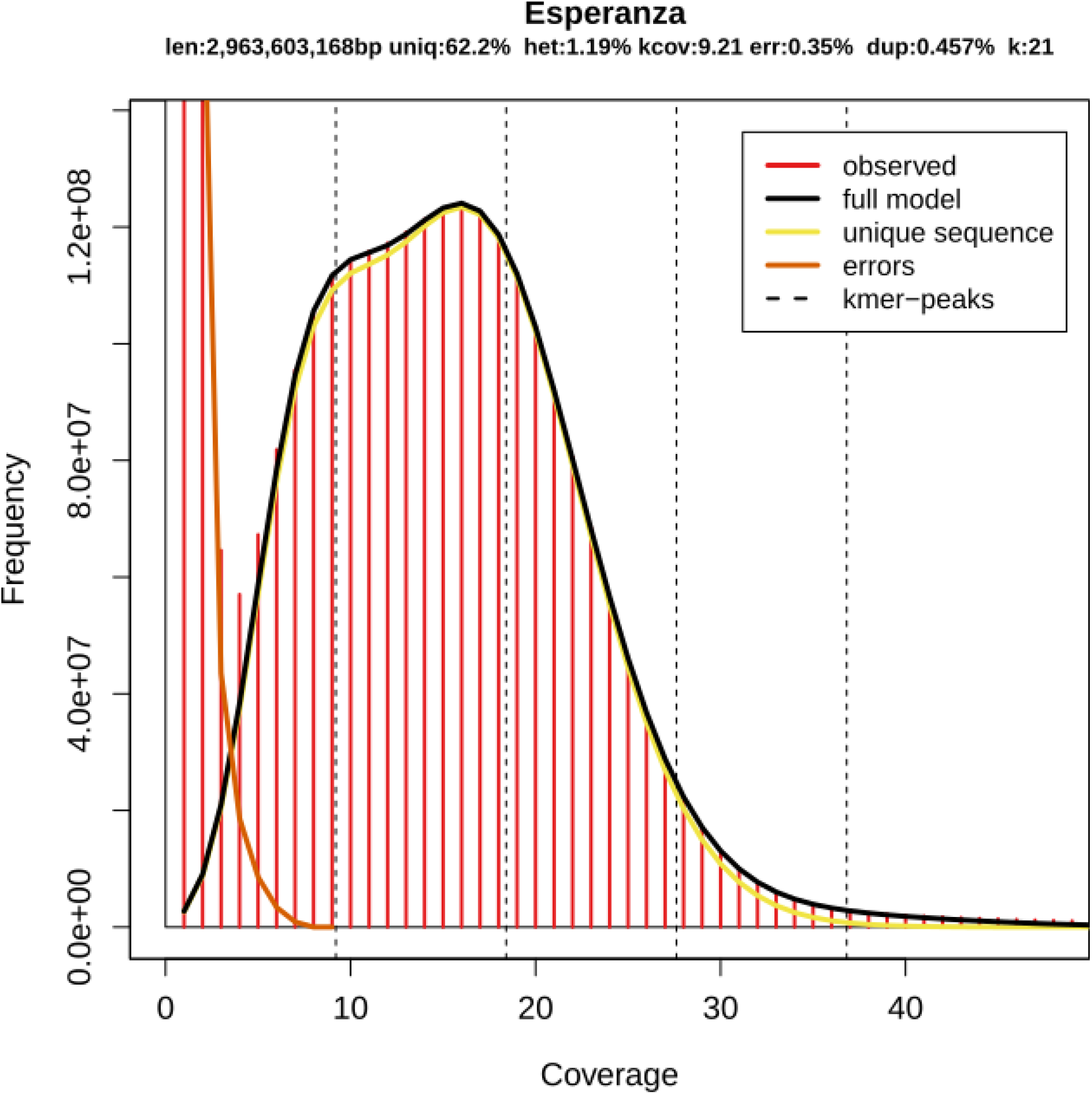
Histogram of 21-mer coverage in short reads from Esperanza gives a genome heterozygosity estimate of ~1.2%.

**Figure S2.**
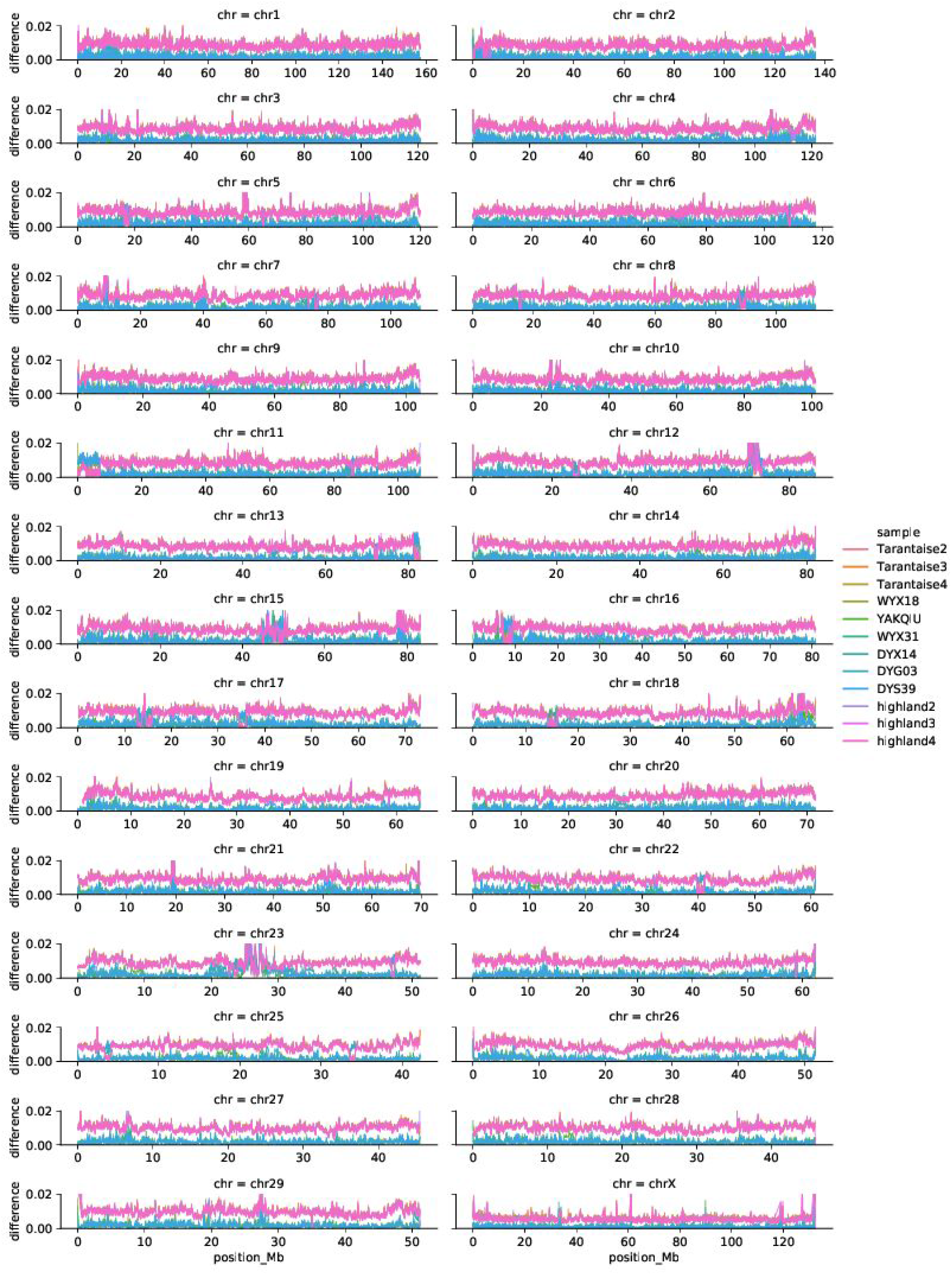
Comparison of twelve yak and cattle genomes to maternal haplotype assembly.

**Figure S3.**
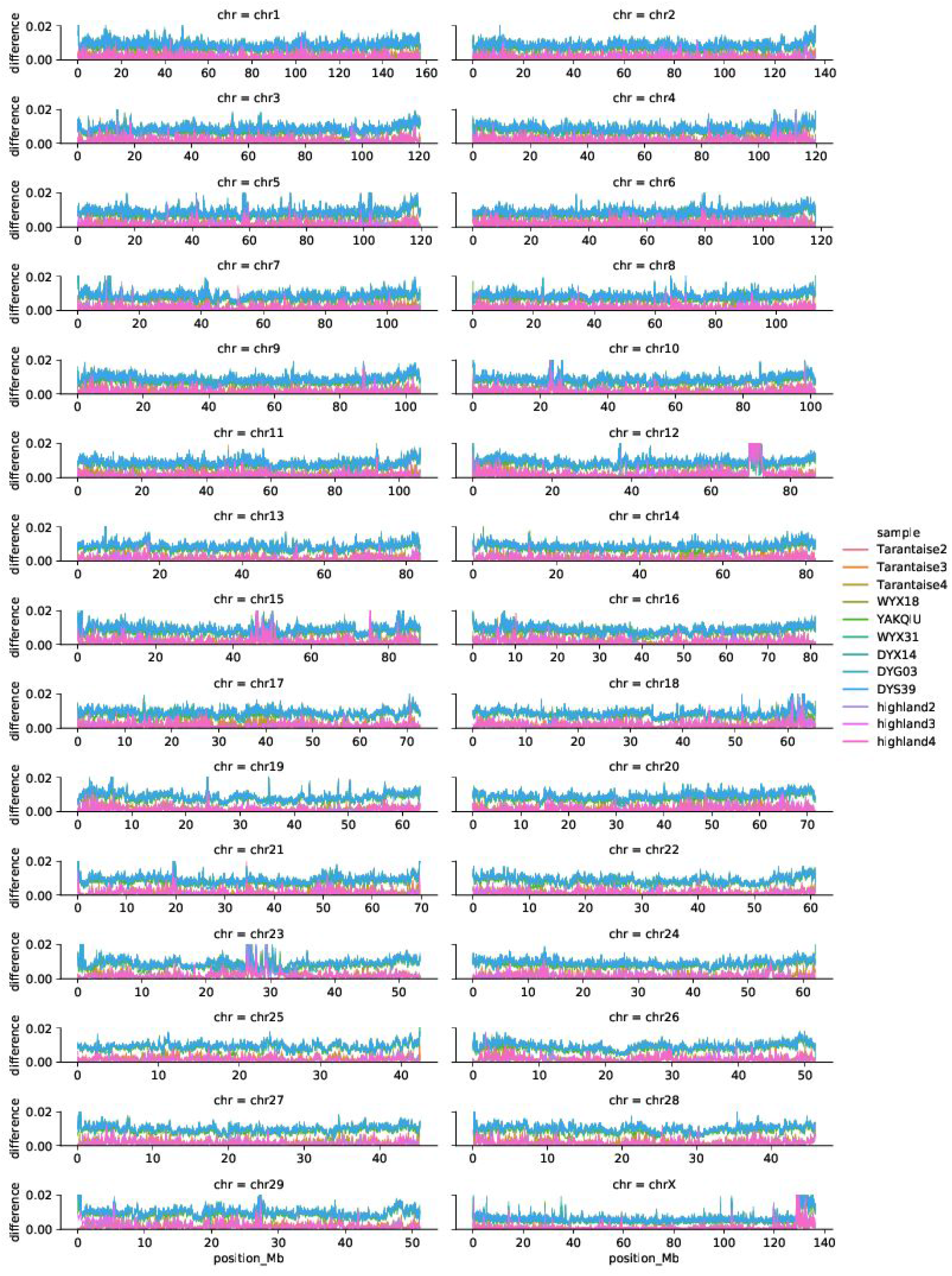
Comparison of twelve yak and cattle genomes to paternal haplotype assembly.

**Supplementary Figure S4.**
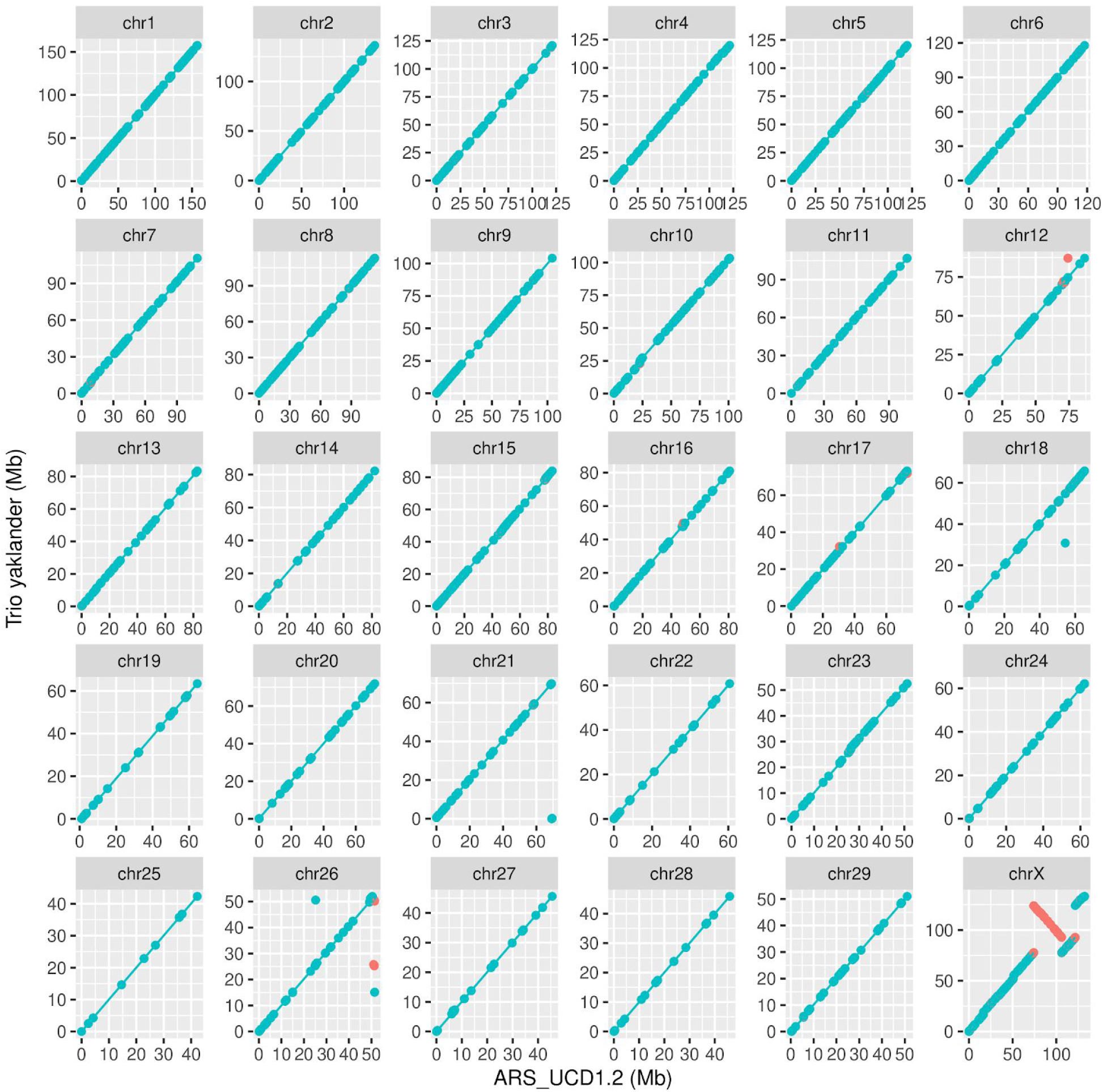
Dot plots of alignment of maternal/yak assembly vs. ARS_UCD1.2, the current cattle reference genome, by chromosome. Blue and red colors denote forward and reverse matches, respectively.

**Supplementary Figure S5.**
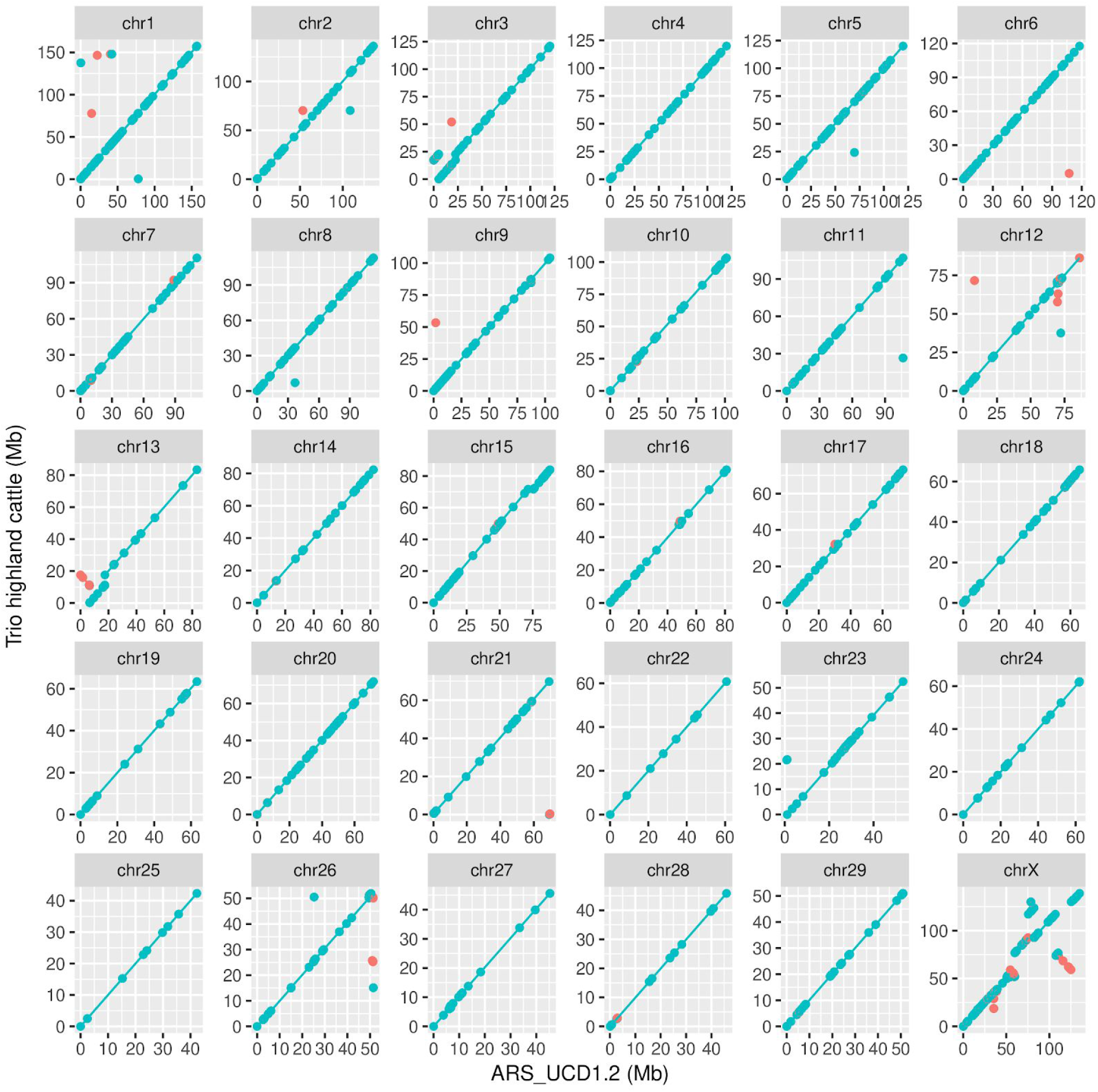
Dot plots of alignment of paternal/highland cattle assembly vs. ARS_UCD1.2, the current cattle reference genome, by chromosome. Blue and red colors denote forward and reverse matches, respectively.

## Supplementary Tables

**Supplementary Table S1 (attached):** Results of verification of Esperanza’s parentage.

**Supplementary Table S2 (attached):** SRA accessions of yak and cattle libraries used for phasing confirmation.

**Supplementary Table S3:** Intersection of repetitive elements with gaps

|  | Sire |  | Dam |  |
| --- | --- | --- | --- | --- |
| Repeat Status | Closed | Trans-Scaffold | Closed | Trans-Scaffold |
| Repeat Consistent | 35 | 25 | 29 | 25 |
| Repeat Complex | 175 | 142 | 172 | 152 |
| No Repeat | 9 | 12 | 12 | 8 |

**Supplementary Table S4:** Frequency of repetitive elements flanking gap regions

| Repeat Class | Count |
| --- | --- |
| Line/L1 | 391 |
| Line/RTE-BovB | 288 |
| Sine/tRNA-Core-RTE | 167 |
| Sine(other) | 223 |
| Simple Repeat | 118 |
| LTR-ERV1 | 72 |
| Satellite/Centromere | 65 |

**Supplementary Table S5:** Consistency of gap closures between sire and dam assemblies

|  | Consistent Gap status | Not Consistent |
| --- | --- | --- |
| Closed | 110 | 109 |
| Trans-Scaffold | 102 | 77<sup>1</sup> |

<sup>1</sup> These discrepancies indicate a structural deviation between the assemblies that may be sub-species specific.

## References

1. Rice ES, Green RE. New Approaches for Genome Assembly and Scaffolding. Annu Rev Anim Biosci [Internet]. 2018; Available from: http://dx.doi.org/10.1146/annurev-animal-020518-115344

2. Alkan C, Sajjadian S, Eichler EE. Limitations of next-generation genome sequence assembly. Nat Methods. Nature Research; 2011;8:61–5.

3. Treangen TJ, Salzberg SL. Repetitive DNA and next-generation sequencing: computational challenges and solutions. Nat Rev Genet. NIH Public Access; 2012;13:36–46.

4. Ardui S, Ameur A, Vermeesch JR, Hestand MS. Single molecule real-time (SMRT) sequencing comes of age: applications and utilities for medical diagnostics. Nucleic Acids Res. 2018;46:2159–68.

5. Payne A, Holmes N, Rakyan V, Loose M. Whale watching with BulkVis: A graphical viewer for Oxford Nanopore bulk fast5 files [Internet]. bioRxiv. 2018 [cited 2019 Mar 8]. p. 312256. Available from: https://www.biorxiv.org/content/10.1101/312256v1.abstract.

6. Jain M, Koren S, Miga KH, Quick J, Rand AC, Sasani TA, et al. Nanopore sequencing and assembly of a human genome with ultra-long reads. Nat Biotechnol. 2018;36:338–45.

7. Low WY, Tearle R, Bickhart DM, Rosen BD, Kingan SB, Swale T, et al. Chromosome-level assembly of the water buffalo genome surpasses human and goat genomes in sequence contiguity. Nat Commun. 2019;10:260.

8. Koren S, Walenz BP, Berlin K, Miller JR, Bergman NH, Phillippy AM. Canu: scalable and accurate long-read assembly via adaptive k-mer weighting and repeat separation. Genome Res. 2017;27:722–36.

9. Kolmogorov M, Yuan J, Lin Y, Pevzner P. Assembly of long error-prone reads using repeat graphs [Internet]. bioRxiv. 2018 [cited 2018 Apr 5]. p. 247148. Available from: https://www.biorxiv.org/content/early/2018/01/12/247148.

10. Li H. Minimap and miniasm: fast mapping and de novo assembly for noisy long sequences. Bioinformatics.2016;32:2103–10.

11. Chin C-S, Peluso P, Sedlazeck FJ, Nattestad M, Concepcion GT, Clum A, et al. Phased diploid genome assembly with single-molecule real-time sequencing. Nat Methods. 2016;13:1050–4.

12. Kronenberg ZN, Hall RJ, Hiendleder S, Smith TPL, Sullivan ST, Williams JL, et al. FALCON-Phase: Integrating PacBio and Hi-C data for phased diploid genomes [Internet]. bioRxiv. 2018 [cited 2019 Mar 8]. p. 327064. Available from: https://www.biorxiv.org/content/10.1101/327064v1

13. Weisenfeld NI, Kumar V, Shah P, Church DM, Jaffe DB. Direct determination of diploid genome sequences. Genome Res. 2017;27:757–67.

14. Koren S, Rhie A, Walenz BP, Dilthey AT, Bickhart DM, Kingan SB, et al. De novo assembly of haplotype-resolved genomes with trio binning. Nat Biotechnol [Internet]. 2018; Available from: http://dx.doi.org/10.1038/nbt.4277.

15. Qiu Q, Zhang G, Ma T, Qian W, Wang J, Ye Z, et al. The yak genome and adaptation to life at high altitude. Nat Genet. 2012;44:946–9.

16. Tumennasan K, Tuya T, Hotta Y, Takase H, Speed RM, Chandley AC. Fertility investigations in the F_1_ hybrid and backcross progeny of cattle (*Bos taurus*) and yak (*B. grunniens*) in Mongolia. Cytogenet Cell Genet. 1997;78:69–73.

17. Simão FA, Waterhouse RM, Ioannidis P, Kriventseva EV, Zdobnov EM. BUSCO: assessing genome assembly and annotation completeness with single-copy orthologs. Bioinformatics. Oxford University Press; 2015;31:3210–2.

18. Medugorac I, Graf A, Grohs C, Rothammer S, Zagdsuren Y, Gladyr E, et al. Whole-genome analysis of introgressive hybridization and characterization of the bovine legacy of Mongolian yaks. Nat Genet. nature.com; 2017;49:470–5.

19. Wu D-D, Ding X-D, Wang S, Wójcik JM, Zhang Y, Tokarska M, et al. Pervasive introgression facilitated domestication and adaptation in the Bos species complex. Nat Ecol Evol. 2018;2:1139–45.

20. Ma L, O’Connell JR, VanRaden PM, Shen B, Padhi A, Sun C, et al. Cattle Sex-Specific Recombination and Genetic Control from a Large Pedigree Analysis. PLoS Genet. 2015;11:e1005387.

21. Zhou S, Goldstein S, Place M, Bechner M, Patino D, Potamousis K, et al. A clone-free, single molecule map of the domestic cow (Bos taurus) genome. BMC Genomics. 2015;16:644.

22. Bovine Genome Sequencing and Analysis Consortium, Elsik CG, Tellam RL, Worley KC, Gibbs RA, Muzny DM, et al. The genome sequence of taurine cattle: a window to ruminant biology and evolution. Science. science.sciencemag.org; 2009;324:522–8.

23. Behl JD, Verma NK, Tyagi N, Mishra P, Behl R, Joshi BK. The major histocompatibility complex in bovines: a review. ISRN Vet Sci. 2012;2012:872710.

24. Takeshima S-N, Sasaki S, Meripet P, Sugimoto Y, Aida Y. Single nucleotide polymorphisms in the bovine MHC region of Japanese Black cattle are associated with bovine leukemia virus proviral load. Retrovirology. 2017;14:24.

25. Fontanesi L, Tazzoli M, Russo V, Beever J. Genetic heterogeneity at the bovine KIT gene in cattle breeds carrying different putative alleles at the spotting locus. Anim Genet. 2010;41:295–303.

26. Tazzoli M, Beever JE, Fontanesi L, Russo V. Identification of mutations in the bovine KIT gene, a candidate for the Spotted locus in cattle. Ital J Anim Sci. Taylor & Francis; 2007;6:218–218.

27. ARS-UCD1.2 - Genome - Assembly - NCBI [Internet]. [cited 2019 Mar 12]. Available from:https://www.ncbi.nlm.nih.gov/assembly/GCF_002263795.1/

28. Bickhart DM, Rosen BD, Koren S, Sayre BL, Hastie AR, Chan S, et al. Single-molecule sequencing and chromatin conformation capture enable de novo reference assembly of the domestic goat genome. Nat Genet. Nature Publishing Group; 2017;49:643–50.

29. Kalbfleisch TS, Rice ES, DePriest MS Jr, Walenz BP, Hestand MS, Vermeesch JR, et al. Improved reference genome for the domestic horse increases assembly contiguity and composition. Commun Biol. 2018;1:197.

30. Jain M, Olsen HE, Turner DJ, Stoddart D, Bulazel KV, Paten B, et al. Linear assembly of a human centromere on the Y chromosome. Nat Biotechnol. 2018;36:321–3.

31. Heaton MP, Keele JW, Harhay GP, Richt JA, Koohmaraie M, Wheeler TL, et al. Prevalence of the prion protein gene E211K variant in U.S. cattle. BMC Vet Res. 2008;4:25.

32. Vurture GW, Sedlazeck FJ, Nattestad M, Underwood CJ, Fang H, Gurtowski J, et al. GenomeScope: fast reference-free genome profiling from short reads. Bioinformatics. 2017;33:2202–4.

33. Heaton MP, Smith TPL., Carnahan JK, Basnayake V, Qiu J, Simpson B, et al. Using diverse U.S. beef cattle genomes to identify missense mutations in EPAS1, a gene associated with pulmonary hypertension. F1000Res. 2016;5:2003.

34. Heaton MP, Smith TPL., Freking BA, Workman AM, Bennett GL, Carnahan JK, et al. Using sheep genomes from diverse U.S. breeds to identify missense variants in genes affecting fecundity. F1000Res. 2017;6:1303.

35. DePristo MA, Banks E, Poplin R, Garimella KV, Maguire JR, Hartl C, et al. A framework for variation discovery and genotyping using next-generation DNA sequencing data. Nat Genet.. 2011;43:491–8.

36. Li H. Aligning sequence reads, clone sequences and assembly contigs with BWA-MEM [Internet]. arXiv [q-bio.GN]. 2013. p. 1303.3997v2. Available from: http://arxiv.org/abs/1303.3997v2

37. Li H, Handsaker B, Wysoker A, Fennell T, Ruan J, Homer N, et al. The Sequence Alignment/Map format and SAMtools. Bioinformatics. 2009;25:2078–9.

38. Ghurye J, Rhie A, Walenz BP, Schmitt A, Selvaraj S, Pop M, et al. Integrating Hi-C links with assembly graphs for chromosome-scale assembly. bioRxiv. Cold Spring Harbor Laboratory; 2018;261149.

39. Jain C, Dilthey A, Koren S, Aluru S, Phillippy AM. A fast approximate algorithm for mapping long reads to large reference databases [Internet]. bioRxiv. 2017 [cited 2019 Mar 5]. p. 103812. Available from: https://www.biorxiv.org/content/10.1101/103812v2

40. English AC, Richards S, Han Y, Wang M, Vee V, Qu J, et al. Mind the Gap: Upgrading Genomes with Pacific Biosciences RS Long-Read Sequencing Technology. PLoS One. Public Library of Science; 2012;7:e47768.

41. Chaisson MJ, Tesler G. Mapping single molecule sequencing reads using basic local alignment with successive refinement (BLASR): application and theory. BMC Bioinformatics. 2012;13:238.

42. Hagberg A, Swart P, S Chult D. Exploring network structure, dynamics, and function using NetworkX [Internet]. Los Alamos National Lab.(LANL), Los Alamos, NM (United States); 2008. Available from: http://conference.scipy.org/proceedings/SciPy2008/paper_2/full_text.pdf

43. Quinlan AR, Hall IM. BEDTools: a flexible suite of utilities for comparing genomic features. Bioinformatics. 2010;26:841–2.

